# Comparative population genomics in two sympatric species of *Strophostyles* (Fabaceae) with contrasting life histories

**DOI:** 10.1101/2021.06.08.447599

**Authors:** Sterling A. Herron, Zachary N. Harris, Matthew J. Rubin, Allison J. Miller

## Abstract

**PREMISE OF THE STUDY:** Life history is an important predictor of population genetic variation, although this link remains unexplored in numerous important plant lineages. One such lineage is the legume genus *Strophostyles*, which contains both annual and herbaceous perennial vines native to eastern North America. Specifically, it remains to be determined whether *Strophostyles* species with different life histories show different patterns of genetic differentiation and diversity, as well as if these species hybridize across their range.

**METHODS:** Here we sampled the perennial *Strophostyles helvola* and annual *S. leiosperma* in five sites across a latitudinal transect in the central United States, including three sites where the species occur in sympatry. Using genotyping-by-sequencing, we identified 5556 polymorphic SNPs across 166 individuals.

**KEY RESULTS:** There is no evidence that *Strophostyles helvola* and *S. leiosperma* hybridize in the populations examined. Within species, *Strophostyles helvola* (perennial) displays admixture among populations, while *S. leiosperma* (annual) does not, although both species show more genetic diversity among rather than within populations. Patterns of genetic diversity are varied across populations of both species, with both heterozygote excess and deficiency.

**CONCLUSIONS:** The complex patterns of genetic differentiation and diversity warrant further investigation of life history and population dynamics in *Strophostyles*, particularly mating system and modes of gene flow. This study exemplifies the diversity of population genetic patterns even within a small genus, and it reinforces the need to characterize such diversity in non-model systems to gain a more complete understanding of how life history contributes to population genetics.

---

Life history strategy has long been recognized as an important predictor of population genetic variation (Loveless and Hamrick, 1984). In general, annual species are expected to have greater population genetic differentiation (*F*_ST_ or *G*_ST_) and lower within-population genetic diversity (*H*_S_), compared to perennial species (Loveless and Hamrick, 1984; Morishima and Barbier, 1990; Austerlitz et al., 2000). However, these genetic associations with life span may be more directly related to other life history traits which are often associated with life span, particularly mating system (selfing, mixed-mating, or outcrossing; Balfourier et al., 1998; Duminil et al., 2007; Duminil et al., 2009). Selfing or mixed-mating species tend to have greater among-population genetic differentiation and lower within-population genetic diversity as compared to outcrossing species (Loveless and Hamrick, 1984; Hamrick and Godt, 1996; Duminil et al., 2007; Gamba and Muchhala, 2020). Often annuals are selfing and perennials are outcrossing, although this pattern is not universal, particularly in herbaceous species (Barrett et al., 1996; Barrett and Harder, 2017). Considering variability over geographic space is also particularly important as a species’ life history may change with differing climates or habitats (Peterson et al., 2016), which can in turn change patterns of population genetic structure (Vest, 2019). Overall, life span alone is not likely to consistently predict population genetic differentiation and diversity, and a broader perspective including other life history traits, the native ecology, and the evolutionary history of the study species and its populations is imperative.

The legume family (Fabaceae Lindl.) is one of the most economically important and widespread groups of plants, containing ca. 19,500 species, and likewise a striking array of life history diversity (Lewis et al., 2005; Azani et al., 2017). Within Fabaceae, subtribe Phaseolinae Bronn (tribe Phaseoleae Bronn ex DC.) stands out, containing the economically important genera *Phaseolus* L. (common bean) and *Vigna* Savi (cowpea), in addition to 25 other wild genera spread throughout Africa, Asia, Australia, and the Americas (Delgado-Salinas et al., 2011). Despite the economic interest of the subtribe, numerous species, particularly those outside genera containing major crops, remain uncharacterized regarding life history and genetic diversity. Assessment of this diversity will be beneficial in understanding the evolutionary history of these lineages and the potential range of genetic variation in crop wild relatives.

*Strophostyles* Elliott (fuzzybean) is a monophyletic legume genus within Phaseolinae, consisting of three species of herbaceous vine: *S. helvola* (L.) Elliott, *S. leiosperma* (Torr. & A. Gray) Piper, and *S. umbellata* (Muhl. ex Willd.) Britton (Riley-Hulting et al., 2004; Delgado-Salinas et al., 2011). Their native range is centered in the eastern United States, extending from Texas east to Florida, north to the Great Lakes and eastern Canada (USDA, 2018; Pelotto and del Pero Martinez, 1998; Riley-Hulting et al., 2004). *S. helvola* is widely distributed across the central and eastern United States, *S. leiosperma* has the westernmost distribution and is primarily restricted to a central band from Minnesota south to Texas, and *S. umbellata* is distributed more in the southeastern portion of the U.S. (Riley-Hulting et al., 2004). *S. helvola* is known to occur in sympatry with *S. leiosperma* in its western range and *S. umbellata* in its eastern range, but *S. leiosperma* and *S. umbellata* are not known to co-occur (Riley-Hulting et al., 2004). All *Strophostyles* species inhabit mesic environments, including freshwater swamps and flood basins, coastal saltwater areas, sand prairies, and anthropogenically disturbed sites such as ditches and ponds (Riley-Hulting et al., 2004). *S. leiosperma* is however distinct in that it occurs in relatively more open, drier environments (Riley-Hulting et al., 2004). *Strophostyles* is also one of the few Phaseolinae clades that has a broad temperate distribution, and it is the only genus in Phaseolinae to have a distribution centered in the United States (Riley-Hulting et al., 2004). *Strophostyles* is sister to the genus *Dolichopsis* Hassl., which is native to the Gran Chaco in South America, an area that also undergoes freezing temperatures, and its other closest genera, *Macroptilium* (Benth.) Urb., *Mysanthus* G.P. Lewis & A. Delgado, and *Oryxis* A. Delgado & G.P. Lewis, are also native to the neotropics (Riley-Hulting et al., 2004). *Strophostyles* and *Dolichopsis* share keel morphology, where the keel gradually curves to the right of the flower as opposed to a sharp curling of the keel as in many other New World Phaseolinae, and the keel in both genera is also distinctly gibbous (Riley-Hulting et al., 2004). *Strophostyles* species are morphologically distinguished from closely related Phaseolinae genera by their cellular seed coating (which lends buoyancy), divergent stipules, a calyx consisting of four lobes that are attenuate to acute, and persistent floral bracts on the pedicels (Riley-Hulting et al., 2004; Delgado-Salinas et al., 2011). Within *Strophostyles*, species are primarily diagnosed by their flower’s keel morphology, although *S. leiosperma* in particular possesses other unique characteristics, including distinctly sericeous leaves and pods, and usually smooth seeds (Riley-Hulting et al., 2004). All three *Strophostyles* species are diploid (2n=22; Lackey, 1980; Riley-Hulting et al., 2004; Yatskievych, 2013).

*Strophostyles* species also have potential in agricultural applications. *S. helvola* and *S. leiosperma* have been targeted as a sustainable, native forage and hay supplement for livestock and wildlife in the southern Great Plains region (Muir et al., 2005; Foster et al., 2008), with cultivars of *S. leiosperma* being developed specifically for that purpose (Butler and Muir, 2010). In the wild, *Strophostyles* herbage and seeds are known to be an important food source for wildlife (Bird and Bird, 1931; Wiseman, 1977; Immel, 2001; Gee et al., 2011). Unlike many cultivated Phaseolinae species, *S. helvola* in particular is known for its salinity tolerance in coastal populations (Tsang and Maun, 1999; Zhang et al., 2018; Zuelsdorf, 2018). *Strophostyles* seeds and roots were also historically consumed by indigenous American peoples, highlighting their human palatability (Parker, 1991; Immel, 2001).

Despite its potential utility and botanical interest, various aspects of *Strophostyles* life history remain unclear. Reported life span of *Strophostyles* species is variable in the literature: *Strophostyles helvola* and *S. leiosperma* have both been reported as either annual or perennial (McGregor et al., 1986; Stubbendieck and Conard, 1989; Isely, 1998; Diggs et al., 1999; Yatskievych, 2013). Nevertheless, the most thorough taxonomic and morphological treatment of the genus to date describes *Strophostyles helvola* as a perennial with a large taproot, and *S. leiosperma* as an annual to short-lived perennial with a small taproot, where the taproot functions as the persisting perennial organ after the annual shoot dies back during the winter (Riley-Hulting et al., 2004; Delgado-Salinas, pers. comm. 2021). Balancing immediate reproduction vs. survival (vegetative allocation) is a major trade-off concept of plant life history theory, is a correlate of plant life span, and is likely to vary by environment (Charnov and Schaffer, 1973; Friedman, 2020; Lundgren and Des Marais, 2020). Specifically, intraspecific variation in life span is associated with environmental disturbance, with annual forms associating with seasonally disturbed sites (often drought) where adult survival is unlikely across seasons or disturbance events, and perennial forms are associated with relatively stable sites where adult survival is likely across seasons (Hall and Willis, 2006; Peterson et al., 2016; Monroe et al., 2019).

Mating system is another aspect of life history likely to be an important predictor of genetic diversity patterns in *Strophostyles*. All *Strophostyles* species have overlapping flowering times and attract a range of generalist bee pollinators (particularly *Megachile*; Robertson, 1890; Krombein et al., 1979; Riley-Hulting et al., 2004). *S. helvola* specifically has an observed keel mechanism which serves to retrieve foreign pollen from the floral visitors while also depositing its own pollen on the visitor’s thorax via a style-brush, encouraging outcrossing (Foerste, 1885; Robertson, 1890). The pollen deposition mechanism of the other two species is unclear, but this combined with pollinator observations point to at least a partially outcrossing mating system in *Strophostyles* species. *S. leiosperma* is also hypothesized to have at least a partially selfing mating system, based on the observation that closed flowers still produce viable seeds; reproductive assurance through selfing would also be consistent with the annual life span (Riley-Hulting et al., 2004).

Some phylogenetic and transcriptomic studies have targeted *Strophostyles* and its close relatives, but extensive genomic information is lacking for the genus. Riley-Hulting et al. (2004) used chloroplast and ITS markers in a phylogenetic study to confirm all three *Strophostyles* species as a monophyletic clade, as well as to confirm its phylogenetic placement among a number of neotropical Phaseolinae genera. More recently, Zhang et al. (2018) sequenced the transcriptome of *S. helvola* in order to discover upregulated genes associated with salinity tolerance. However, a whole genome population-level approach has yet to be implemented in *Strophostyles*, which leaves a number of open questions. Specifically, while there is little evidence to date for putative hybridization between species in the genus (Riley-Hulting et al., 2004), this has not been directly investigated in sympatric populations of *Strophostyles* species. Also, while there is preliminary evidence for broad differences in genetic diversity across the range of *Strophostyles* (Riley-Hulting et al., 2004), this has not been shown among multiple individuals within multiple populations. Characterization of population genomics may further reveal signatures of mating system and how patterns of dispersal contribute to gene flow among populations of each *Strophostyles* species.

Here we focus on the sympatric species *Strophostyles helvola* and *S. leiosperma*, in the context of their different life history strategies, in populations across a latitudinal transect from northern Iowa to central Arkansas, USA. We ask the following questions: (1) are *Strophostyles helvola* and *S. leiosperma* genetically distinct, or is there evidence of hybridization between these species when they occur in sympatry; (2) is there genetic differentiation among populations within each species, or is there evidence of admixture among populations; and (3) what are the patterns of genetic diversity within and among populations for each species?

## MATERIALS AND METHODS

### Sample collection

This study focused on two of the three *Strophostyles* species: *S. helvola* and *S. leiosperma*, due to their close relationship and sympatric distribution in the Midwest United States (Riley-Hulting et al., 2004). We collected samples from five sites along a latitudinal transect ranging from northern Iowa to central Arkansas, USA (Fig. 1; Table 1; Appendix S1; see Supplemental Data with this article). Collection sites were targeted based on occurrence records for the species (Global Biodiversity Information Facility; www.gbif.org) and by contacting regional botanists. *Strophostyles umbellata* was not collected due to its rarity in the sites sampled (it has not been observed in this region by the authors); while it is reported to occur in this range, it does not appear to occur sympatrically with *S. helvola* and *S. leiosperma* at least in the populations observed. In general, *S. umbellata* is known to have more scattered individuals and populations of low density, and thus the likelihood of finding substantial populations is lower (Riley-Hulting et al., 2004).

**Table 1.** Summary of population sampling for *Strophostyles helvola* and *S. leiosperma* across sites. *S. leiosperma* was not present at sites SP-MO and AR. Sites are sorted by descending latitude. Lat/long coordinates are the approximate centroid of the primary area of sampling at each site (average of coordinates of all samples), and the elevation is the rounded average among all samples in meters above sea level (masl). Individual sample information is available in Appendix S1.

| Location(s) | Site ID | Lat/long | Elevation<br>(masl) | Species | Individuals<br>sequenced | Voucher<br>Collection No. |
| --- | --- | --- | --- | --- | --- | --- |
| Cedar Hills Sand Prairie, Cedar Falls, Iowa, USA | IA | 42.59764<br>-92.55197 | 284 | *Strophostyles helvola*<br>*Strophostyles leiosperma* | 20<br>20 | SH 3.8; SH 3.28<br>SH 3.10; SH 3.25 |
| Frost Island Conservation Area & Iliniwek Village State Historic Site, Wayland, Missouri, USA | IV-MO<sup>a</sup> | 40.42992<br>-91.55692 | 167 | *Strophostyles helvola*<br>*Strophostyles leiosperma* | 21<br>17 | SH 2.7; SH 2.32<br>SH 2.6; SH 2.31 |
| Shaw Nature Reserve, Gray Summit, Missouri, USA | SNR-MO | 38.48327<br>-90.81267 | 183 | *Strophostyles helvola*<br>*Strophostyles leiosperma* | 21<br>19 | SH 1.30; SH 1.39<br>SH 1.28; SH 1.33 |
| Sand Pond Conservation Area, Neelyville, Missouri, USA | SP-MO | 36.50350<br>-90.59622 | 93 | *Strophostyles helvola* | 19 | SH 4.20; SH 4.28 |
| Burns Park, North Little Rock, Arkansas, USA | AR<sup>b</sup> | 34.79084<br>-92.32575 | 75 | *Strophostyles helvola* | 29 | SH 7.21; SH 7.22 |

<sup>a</sup> For IV-MO, the approximate centroid and elevation of sampling from Illiniwek Village State Historic Site was used for this table, since that is where the majority of samples were collected from, while five samples were collected approximately 3 km from there at Frost Island Conservation Area.
<sup>b</sup> For AR, the approximate centroid and elevation of sampling from Burns Park was used for this table, since that is where the majority of samples were collected from, while four samples were collected approximately 10 km from there in a gravel ditch in Little Rock, Arkansas.

**Figure 1.**
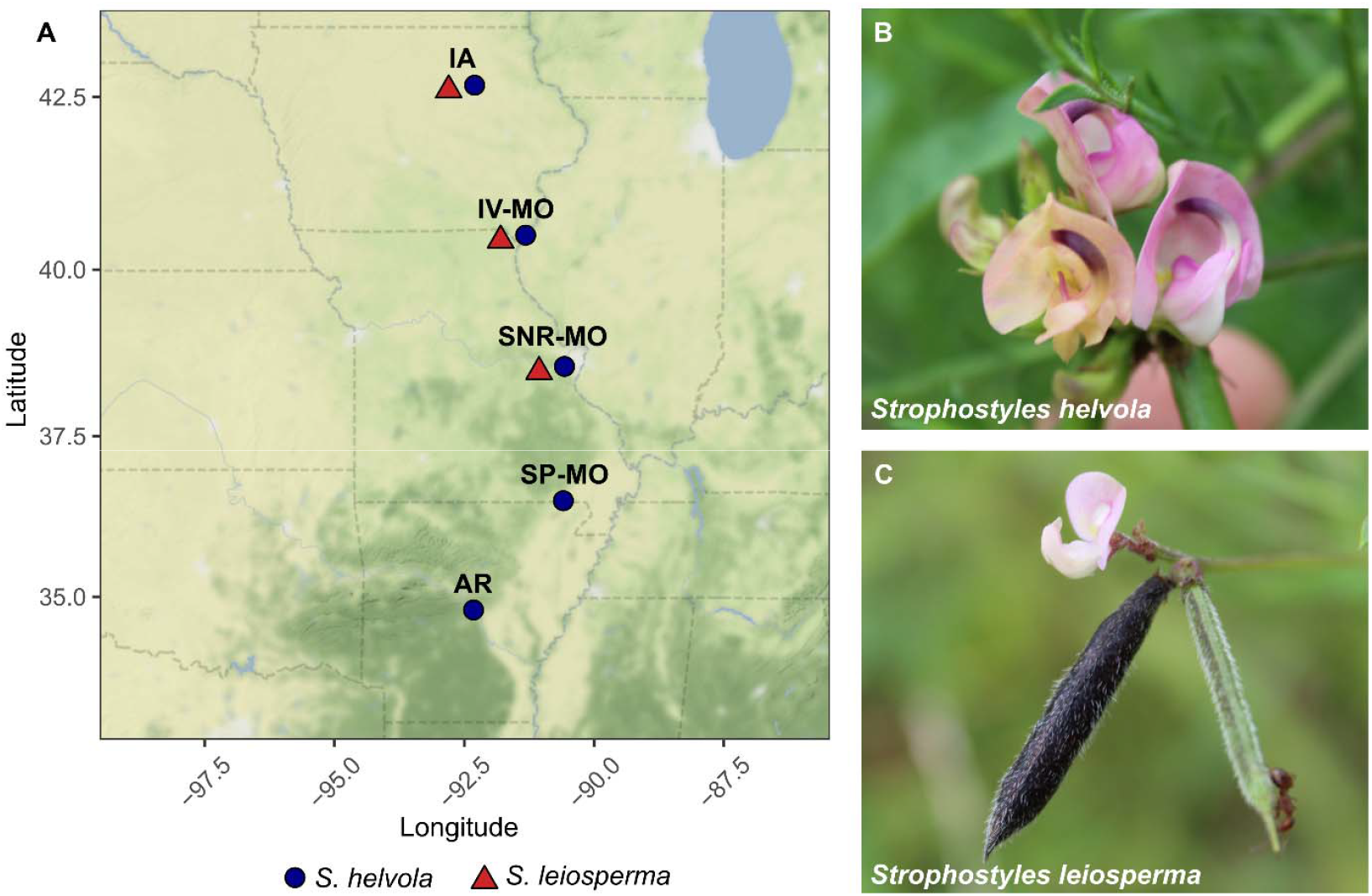
Diagram of population sampling and photographs of *Strophostyles helvola* and *S. leiosperma*. (A) Shows a map of populations sampled from sites across Iowa, Missouri, and Arkansas, USA. *S. helvola* populations are represented by a blue circle and *S. leiosperma* populations are represented by a red triangle. Where the species co-occur at the three northernmost sites, the points are spatially separated for visualization, although they occur at the same coordinates. (B) shows an inflorescence of *S. helvola* at population SNR-MO, and (C) shows an inflorescence of *S. leiosperma* with a flower, unripe pod, and ripe pod, at SNR-MO (not to scale). Note the difference in keel morphology and pubescence on the *S. leiosperma* pod. Photo credit: S.A.H.

From north to south, sites sampled included the Cedar Hills Sand Prairie (Cedar Falls, Iowa, USA; site “IA”), Frost Island Conservation Area and Iliniwek Village State Historic Site (Wayland, Missouri, USA; site “IV-MO”), Shaw Nature Reserve (Gray Summit, Missouri, USA site “SNR-MO”), Sand Pond Conservation Area (Neelyville, Missouri, USA; site “SP-MO”), and Burns Park (North Little Rock, Arkansas, USA; site “AR”) (Fig. 1; Table 1; Appendix S1). Habitat types were generally open prairie, open grassland/shrubland, and forest edges, or the edge of ponds (SNR-MO) and along a riverbank (AR) (Appendix S1). It was confirmed with site managers that *Strophostyles helvola* and *S. leiosperma* were not known to be intentionally seeded at any site (Appendix S1). Linear distance between nearest collection sites ranged from 219 to 256 km, and mean altitude at each site ranged from 76 to 284 meters above sea level (Fig. 1; Table 1). Plant populations at each site were defined as a group of 15 or more individuals of one species within the same 10 square km area. *Strophostyles helvola* and *S. leiosperma* co-occurred in the three northernmost sites (IA, IV-MO, and SNR-MO), while only *S. helvola* was present at SP-MO and AR. At each site, we sampled 17-29 individual plants per species, and in total, we sampled 110 individuals of *S. helvola* and 56 individuals of *S. leiosperma* (Table 1). We collected fresh, young leaf tissue and stored it in paper envelopes, which were sealed in a ziploc bag filled with dry silica for desiccation. Individuals of the same species were collected at least ∼3 m apart. Species were generally identified by their distinct papilionoid flower morphology: *S. helvola* exhibits a prominent and distinctly curved keel, while *S. leiosperma* exhibits a small, only slightly curved keel that is mostly hidden by the wing petals (Fig. 1; Riley-Hulting et al., 2004). In the absence of flowers (rare), these species can also be distinguished on leaf and pod morphology. *S. helvola* has ovate, sparsely haired, often lobed leaflets, while *S. leiosperma* has lanceolate, sericeous, unlobed leaflets (rarely having shallow lobes; Riley-Hulting et al., 2004). *S. helvola* also has large, sparsely haired pods (30-96 mm long) and seeds with an accessory cellular coating, while *S. leiosperma* has smaller, distinctly sericeous pods (12-41 mm long) with smaller, usually smooth seeds (Riley-Hulting et al., 2004). For each plant collected, we recorded the latitude, longitude, and elevation, and photographed most plants to document identifying features. For each population, we also collected two herbarium voucher specimens per species, including the entire aboveground and most of the belowground portion of the plant when possible. We deposited the vouchers at the Missouri Botanical Garden (St. Louis, Missouri, USA; Table 1). The map of population sampling was created using the R packages ‘ggmap’ and ‘ggplot2’ (Kahle and Wickham, 2013; Wickham, 2016; R Core Team, 2021).

### Genomic DNA extraction

We extracted genomic DNA from the leaves of 166 plants collected in the field (Appendix S1). When available, 55-60 mg of dried leaf tissue per sample was ground into powder using liquid nitrogen and a mortar and pestle, or by freezing the tissue in a tube with liquid nitrogen, inserting 0.9-3.175 mm steel ball bearings, and oscillating using a Rech Mixer Mill 400 at 30 Hz for 2 min, or until a powder. DNA was extracted using an E.Z.N.A.^®^ Plant DNA Kit (Omega Bio-tek, Norcross, Georgia, USA) according to the product label, with a few modifications: the initial incubation in P1 buffer was increased to 45-60 min during which samples tubes were vortexed (instead of inverting), and the centrifuge speed and sometimes duration was increased to allow better tissue and DNA pelleting as well as movement of liquid through the DNA mini column. DNA concentration was quantified using a Qubit^®^ dsDNA BR Assay Kit and an Invitrogen™ Qubit™ 3 Fluorometer (Thermo Fisher Scientific, Waltham, Massachusetts, USA). To increase concentration of some samples, we vacufuged them at 30°C and re-eluted them. We retained samples with > 10 ng/μL DNA concentration for sequencing. Eluted DNA was stored at 4°C until shipped for genomic processing (with dry ice).

### Sequencing

We submitted all samples to the University of Minnesota Genomics Center (Minneapolis, Minnesota, USA), where they were sequenced using genotyping-by-sequencing (GBS), a reduced-representation whole genome method (Elshire et al., 2011; Poland et al., 2012). Dual-indexed GBS libraries were created using a double digest with the two enzymes MspI and PstI on 100 ng of DNA. Samples were treated with T4 ligase and phased adaptors with TGCA and CG overhangs. Libraries were pooled and size selected for 300-744 bp segments, and the pool was diluted to 1.5 nM and sequenced on a half-lane of an Illumina NovaSeq 6000 FlowCell with single-end 1×100-bp reads. This generated 54,142 Mb total reads (average 326 Mb per sample), with mean quality score ≥Q30 for all libraries (Appendix S1).

### Alignment and SNP calling

Both ends of sequence reads were trimmed for quality at a minimum Phred score of 30 using bbduk (Bushnell, 2014; qtrim=rl, trimq=30). We aligned quality-filtered reads to the *Phaseolus vulgaris* Andean landrace ‘G19833’ v.2.1 reference genome (https://phytozome-next.jgi.doe.gov/info/Pvulgaris_v2_1), which was downloaded from the Ensembl Genomes website (https://ensemblgenomes.org/; Yates et al., 2020; accessed Feb 26, 2020), using Burrows-Wheeler Alignment (BWA MEM) with default settings (bwa v0.7.17-r1188; Li and Durbin, 2009). In the absence of a *Strophostyles* genome, the *Phaseolus vulgaris* genome was used as a reference genome due to its long history of improvement and high coverage of the genome. Variant calling used bcftools v1.9, with default settings, and genotypes were filtered at a minimum read depth of 10 with vcftools v0.1.16 (--minDP 10; Danecek et al., 2011; all sites passed this filter). Biallelic SNPs were called with PLINK v1.90b4 (Chang et al., 2015). After evaluating different minor allele frequencies and missing genotype rates, a conservative approach was applied: we filtered at a minor allele frequency of 0.025 and a maximum missing genotype rate per SNP of 10% (--biallelic-only --snps-only --maf 0.025 --geno 0.1). Less conservative filtering approaches increased the number of SNPs retained but did not change the broad genetic structure trends or the conclusions. Ultimately, 1,584,797 SNPs were called in total. 5556 SNPs and all 166 samples were retained following filtering.

### Population genetic differentiation and diversity

We implemented fastSTRUCTURE (Raj et al., 2014) to characterize the population genetic structure of all samples, testing values of K clusters ranging from 1 to 8 (total population number considering both species). We then used the chooseK.py function in fastSTRUCTURE to determine the optimal K value based on two criteria: (1) model complexity which maximized marginal likelihood, and (2) the number of model components used to explain structure in the data. K=6 clusters was optimal based on the number of model components, while K=7 was optimal based on maximized marginal likelihood. The seventh cluster group for the K=7 model has an extremely rare representation such that effectively only 6 clusters emerge, so we defer to the original K=6 model for further discussion. Population structure plots were generated using the R packages ‘ggplot2’ and ‘pophelper’ (Francis, 2017). We also implemented principal component analysis (PCA) of genetic variation using PLINK (--pca 166) and visualized the PCA using ‘ggplot2’ with the EIGENVEC output from PLINK.

For calculations of population statistics, our filtered SNP variant call format file was converted to a GENEPOP text file using PGDSpider2 (Lischer and Excoffier, 2012), which was then converted to a GENIND object in R (package ‘adegenet’; Jombart, 2008) and reformatted to a GENEPOP object (‘graph4lg’ package; Savary, 2020) usable in Genodive v.3.04 (Meirmans, 2020), with the modification of changing missing values to ‘0000’ in order to be interpreted by Genodive as diploid. All further analyses were completed in Genodive. Pairwise *F*_ST_ values between all populations within species were calculated and *P*-values generated from 1000 permutations. Isolation by distance was also tested for each species using a Mantel test. Briefly, the test assesses correlation between the geographic distance matrix (based on the average of all sample coordinates for each population of each species, respectively) and the pairwise genetic distance matrix [*F*_ST_/(1-*F*_ST_)] between populations within a species, with 1000 permutations. Genetic diversity metrics were also calculated per population for each species, including the observed heterozygosity (*H*_O_), within-population expected heterozygosity (*H*_S_; Nei’s gene diversity; Nei, 1987), and inbreeding coefficient (*F*_IS_). The deviation of genotype frequencies within each population from the expectation under Hardy-Weinberg equilibrium (random mating) was calculated using the least squares method (from analysis of molecular variance, AMOVA; Excoffier et al., 1992) with 1000 permutations. An AMOVA was also calculated using the Infinite Allele Model with 1000 permutations, in order to dissect the percentage of genetic variance within individuals, among individuals nested within populations, among populations nested within species, and among species.

## RESULTS

### Strophostyles helvola *and* S. leiosperma *are distinct lineages*

*Strophostyles helvola* and *S. leiosperma* were found to be genetically distinct from each other within and across collection sites. *Strophostyles helvola* and *S. leiosperma* separated distinctly with K=2 genetic clusters (Fig. 2). Principal component analysis also supported the species separation: PC1, which separates *S. helvola* and *S. leiosperma*, explained 45% of the total genetic variation (Fig. 3A). Lastly, the analysis of molecular variance (AMOVA) corroborated this pattern, in that a significant percentage of genetic variation was attributed to among-species variation (62.4%; *F*_CT_ = 0.642, *P* = 0.001; Table 2).

**Table 2.** Analysis of molecular variance (AMOVA) results from eight populations with 5556 total SNPS for both *Strophostyles helvola* and *S. leiosperma*. Here we decompose the genetic variation for individuals nested within populations, populations nested within species, among species, and among individuals. *P*-values are derived from 1000 permutations and thus have a lower threshold of 0.001. The unnested *F*_IT_ terms is not interpretable and thus not assigned a *P*-value.

| Source of variation | Nested in | % Variation | *F*-stat | *F*-value | *P*-value |
| --- | --- | --- | --- | --- | --- |
| Among individuals | Population | -0.1 <sup>a</sup> | $F_{IS}$ | -0.003 <sup>a</sup> | 0.937 |
| Among populations | Species | 17.7 | $F_{SC}$ | 0.471 | 0.001 |
| Among species | --- | 62.4 | $F_{CT}$ | 0.624 | 0.001 |
| Among individuals | --- | 19.9 | $F_{IT}$ | 0.801 | --- |

<sup>a</sup> Slightly negative values for percent variation and *F* statistics can occur due to a lack of significant genetic structure; since our results show a lack of significance for this source of variation, we interpret these values as equivalent to zero (Schneider et al. 2000; Meirmans 2006).

**Figure 2.**
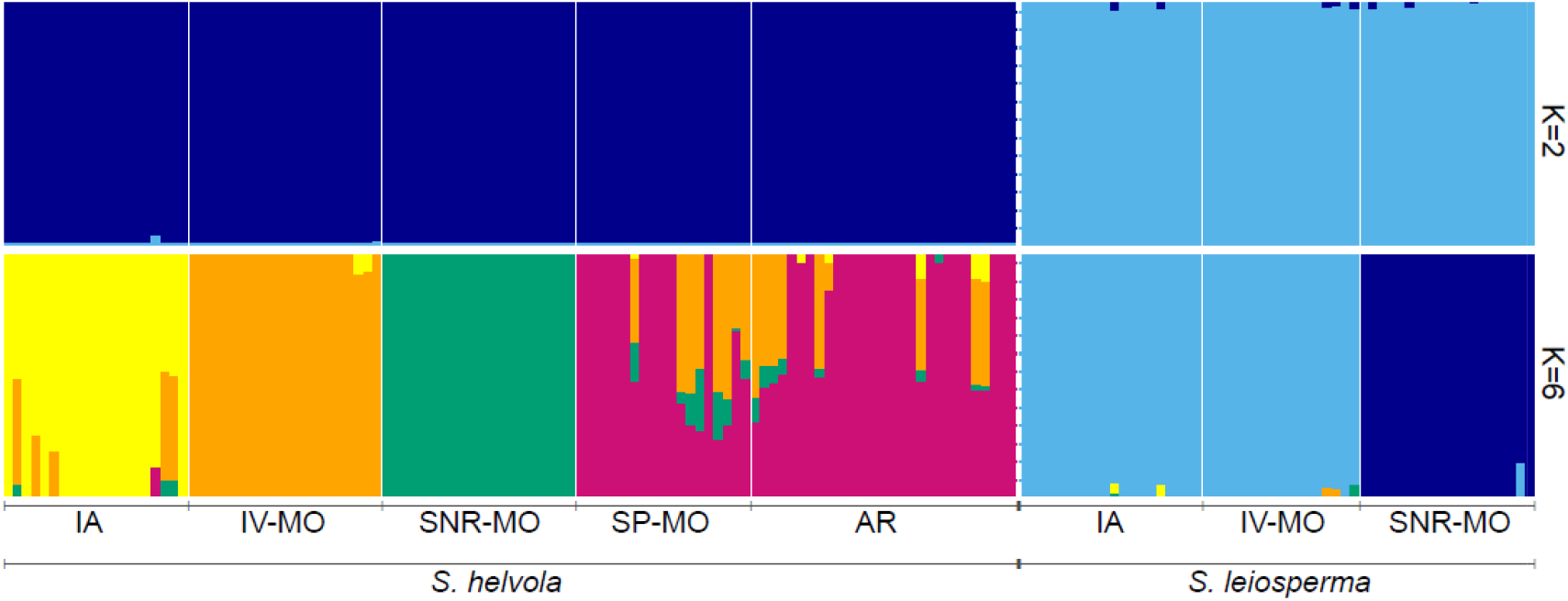
Genetic structure plots with K=2 and K=6 clusters for *Strophostyles helvola* and *S. leiosperma*; K=2 separated the species, and K=6 best explained overall genetic structure in the data. Populations, labeled below the bars, are ordered north to south within species (left to right) and are separated by thin white solid lines; species are separated by a thick white dashed line. Each vertical bar represents a single individual from each population. Each color signifies a unique genetic cluster assignment, with multiple colors within an individual signifying mixed ancestry.

**Figure 3.**
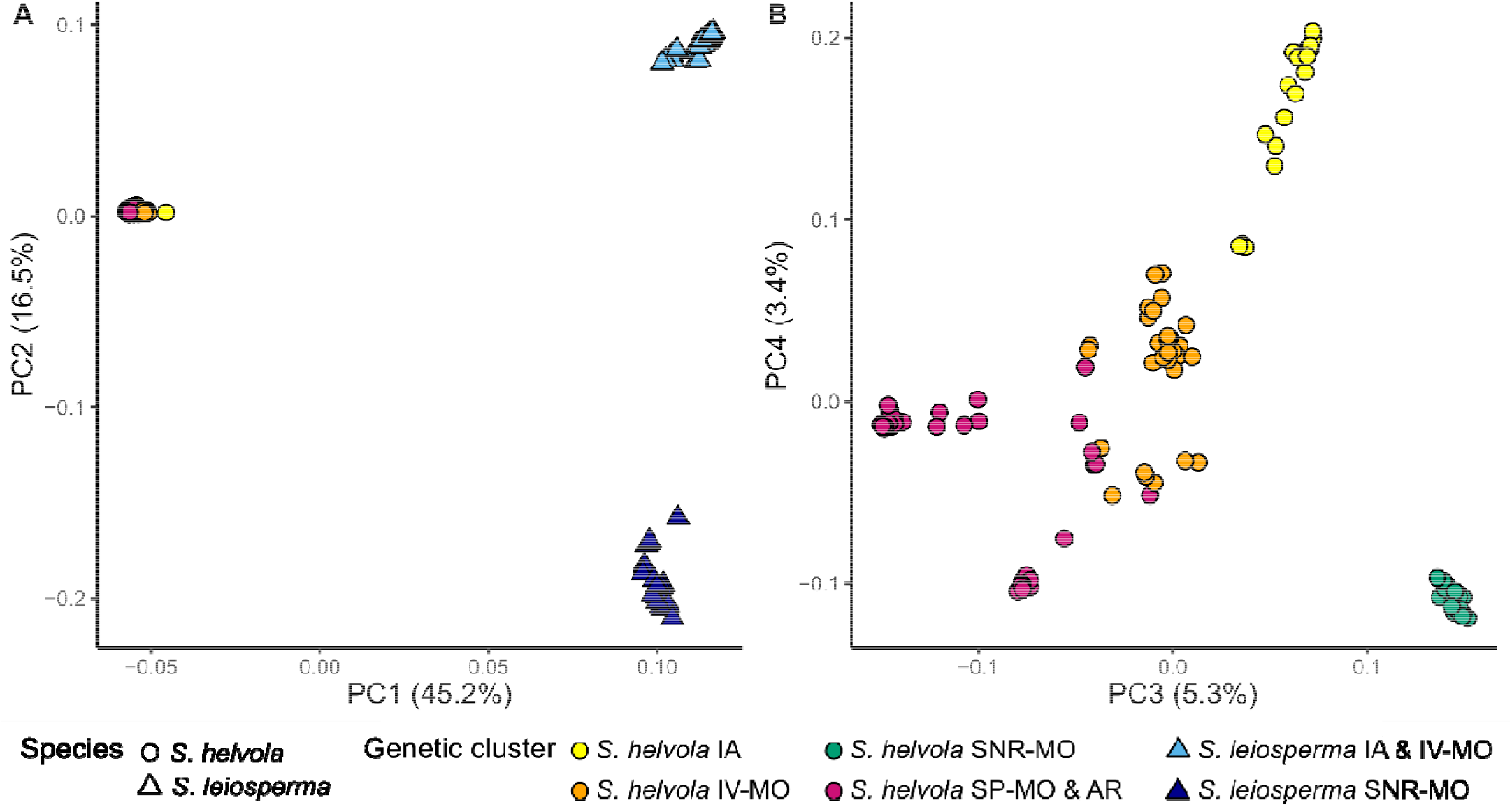
Principal component analysis of SNP data for all populations of *Strophostyles helvola* and *S. leiosperma*, where each point represents a single individual; point fill color matches the predominant genetic cluster assignment from Fig. 2 and point shape corresponds to species. Displayed are (A) PC1 and PC2 (both species), and (B) PC3 and PC4 (just *S. helvola*). *S. leiosperma* is omitted from panel (B) due to nonsignificant genetic variation among populations for PC3 and PC4, and in order to better visualize *S. helvola* variation.

### Within-species population genetic structure differs for Strophostyles helvola and S. leiosperma

There were unique patterns of genetic clustering within each species, with *Strophostyles helvola* showing evidence of admixture among some populations and *S. leiosperma* showing genetically distinct clusters. For K=6 (optimal K value that explained structure in the data, based on the number of model components), individuals of *S. helvola* assigned to one or more of four clusters, and individuals of *S. leiosperma* assigned to two other distinct clusters (Fig. 2). Within *S. helvola*, individuals from populations IV-MO and SNR-MO showed the strongest population differentiation from other clusters (Fig. 2). SNR-MO was the only population of *S. helvola* with no evidence of admixture, while two individuals of *S. helvola* population IV-MO showed only slight admixture with the IA population (Fig. 2). In contrast, *S. helvola* populations IA, SP-MO, and AR showed more evidence of admixture (Fig. 2). For *S. helvola* population IA, 14/20 individuals’ genetic composition assigned only to its own unique genetic cluster, while 6/20 individuals showed admixture, primarily with population IV-MO, but also with SNR-MO and SP-MO/AR (Fig. 2). Most *S. helvola* individuals from population SP-MO and AR have genetic ancestry that assigned predominantly to the same genetic cluster, while the admixture present in both populations assigned primarily to IV-MO (Fig. 2). For population SP-MO specifically, 11/19 individuals assigned only to their own genetic cluster, while the other 8/19 individuals showed signs of admixture, primarily with population IV-MO but also with SNR-MO (Fig. 2). For population AR, 18/29 individuals assigned solely to their own genetic cluster (the same as SP-MO), with the remaining 11/29 again showing admixture mainly with population IV-MO but also IA and SNR-MO (Fig. 2). For *S. leiosperma*, only two genetic clusters emerged and there was a distinct lack of admixture. Both the IA and IV-MO populations assigned to the same genetic cluster, with no substantial evidence of shared ancestry with other clusters (Fig. 2). All 37 individuals of the *S. leiosperma* IA and IV-MO populations assigned predominately to their own genetic cluster (Fig. 2). For *S. leiosperma* population SNR-MO, 18/19 individuals assigned solely to their own genetic cluster, with only one individual showing slight admixture with the other *S. leiosperma* cluster (Fig. 2). Considering intermediate K values between K=2 and K=6, K=3 through K=5 illustrated the stepwise separation of north and south *S. helvola* populations, the two clusters of *S. leiosperma*, and potential mixed ancestry in population IV-MO (Appendix S2). K=7 cluster structure (see Methods) was similar to the K=6 pattern, with the main deviations being that *S. helvola* populations IV-MO and SP-MO primarily assigned to the same genetic cluster, and that population AR primarily assigned to a different genetic cluster than SP-MO (Appendix S2).

Fixation index results largely supported the fastSTRUCTURE patterns. *Strophostyles helvola* population SNR-MO showed some of the highest pairwise *F*_ST_ values with the other *S. helvola* populations (0.359 - 0.468) (Table 3). The strong admixture signal of *S. helvola* population IV-MO with IA, SP-MO, and AR was reflected by its lower *F*_ST_ values when these populations were paired (0.260-0.284) (Table 3). The same predominant genetic cluster assignment of *S. helvola* populations SP-MO and AR was also reflected by their low pairwise *F*_ST_ (0.294) (Table 3). Consistent with its low levels of admixture, *S. helvola* population IA showed high *F*_ST_ values paired with SP-MO (0.383) and AR (0.405) (Table 3). Additionally, *Strophostyles leiosperma* cluster separation was reflected in *F*_ST_ values, where populations IA and IV-MO had a low pairwise *F*_ST_ of 0.251, while the *F*_ST_ of *S. leiosperma* population SNR-MO with IA and IV-MO was 0.712 and 0.707, respectively (Table 3). Based on the Mantel test, we also found no evidence of isolation by distance for either species. There was only a slight, nonsignificant positive correlation between genetic [*F*_ST_/(1-*F*_ST_)] and geographic distance for *S. helvola* (*R*^2^=0.020; *P*=0.362) and *S. leiosperma* (*R*^2^=0.184; *P*=0.495). This is consistent with the nonlinear pattern of genetic similarity across geographic space observed in our fastSTRUCTURE and *F*_ST_ results (Fig. 2; Table 3).

**Table 3.**
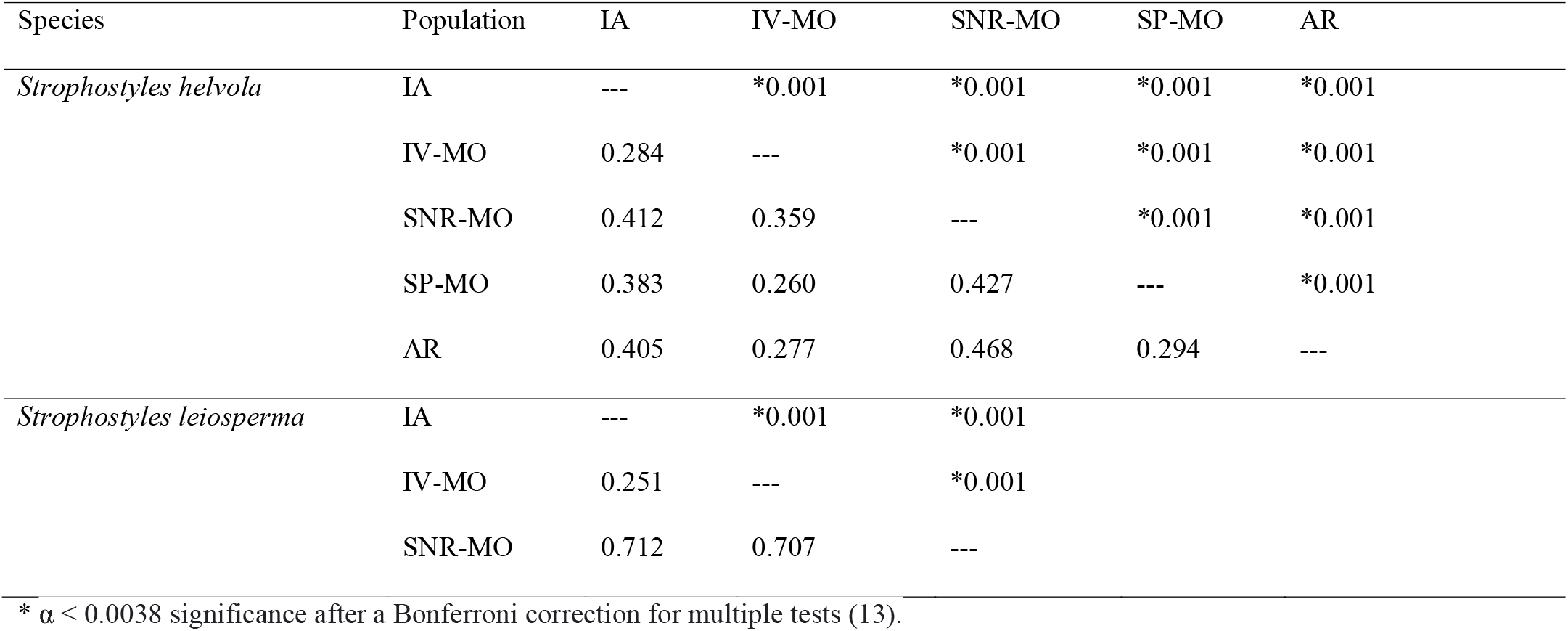
Pairwise *F*_ST_ values between all combinations of populations within species, including five populations of *Strophostyles helvola*, and three populations of *S. leiosperma*, based on 5556 biallelic SNPs. Populations are arranged from north to south latitudes, first with *S. helvola*, then *S. leiosperma. F*_ST_ values are shown below the diagonal, and the *P*-values for the pairwise *F*_ST_ are above the diagonal. *P*-values are derived from 1000 permutations and thus have a lower threshold of 0.001.

| Species | Population | IA | IV-MO | SNR-MO | SP-MO | AR |
| --- | --- | --- | --- | --- | --- | --- |
| *Strophostyles helvola* | IA | --- | *0.001 | *0.001 | *0.001 | *0.001 |
|  | IV-MO | 0.284 | --- | *0.001 | *0.001 | *0.001 |
|  | SNR-MO | 0.412 | 0.359 | --- | *0.001 | *0.001 |
|  | SP-MO | 0.383 | 0.260 | 0.427 | --- | *0.001 |
|  | AR | 0.405 | 0.277 | 0.468 | 0.294 | --- |
| *Strophostyles leiosperma* | IA | --- | *0.001 | *0.001 |  |  |
|  | IV-MO | 0.251 | --- | *0.001 |  |  |
|  | SNR-MO | 0.712 | 0.707 | --- |  |  |

\* $\alpha < 0.0038$ significance after a Bonferroni correction for multiple tests (13).

Principal component analysis also largely supported the intraspecific genetic patterns observed in fastSTRUCTURE and *F*_ST_ values. PC2, which separated the two clusters of *S. leiosperma*, explained 17% of the total genetic variation (Fig. 3A). *S. helvola* populations remained tightly clustered on both PC1 and PC2, with intraspecific variation only emerging in PC3 and PC4 (Fig. 3B). *S. helvola* individuals with admixture from the AR / SP-MO and IV-MO clusters were intermediate to the non-admixed individuals from these clusters in PC3 and PC4 (Fig. 3B). There was also some overlap of the *S. helvola* SNR-MO cluster with the *S. helvola* AR / SP-MO cluster on PC4 alone, reflecting the slight admixture of SNR-MO in those populations (Fig. 3B). The *S. helvola* IA cluster remained relatively isolated on PC3 and PC4, with the exception of a few admixed individuals approaching the *S. helvola* IV-MO cluster, representing the admixture between those populations (Fig. 3B). *S. leiosperma* clusters all lacked genetic variation on PC3 (*F*_1_=3.11; *P*=0.08) and PC4 (*F*_1_=1.61; *P*=0.21) and thus were omitted from Fig. 3B in order to better visualize variation in *S. helvola*.

The AMOVA further confirmed that, in addition to genetic variation between species, there was a significant level of variation among populations within species (17.7%; *F*_SC_=0.471; *P*=0.001), but a lack of significant variation among individuals within populations (−0.1%; *F*_IS_=-0.003; *P*=0.937; Table 2; slight negative deviations of *F* statistics from 0 can be interpreted as a lack of genetic structure among members of that group, i.e., *F*_IS_≈0; Schneider et al., 2000; Meirmans, 2006).

### Population genetic diversity patterns are varied

Both species showed mixed patterns of genetic diversity among their populations. The genotype frequencies of all populations deviated significantly from Hardy-Weinberg equilibrium (*P* = 0.001-0.003) with the exception of *S. leiosperma* population SNR-MO (*P* = 0.487), both due to heterozygote excess and deficiency as determined by *F*_IS_ (Table 4). The greatest observed heterozygosity (*H*_O_) was observed in the *S. helvola* IA population (0.109) and the lowest in the *S. helvola* SNR-MO population (0.087; Table 3). *S. leiosperma* had more consistently high *H*_O_ across its populations (0.100-0.108) than *S. helvola* (Table 4). In contrast, *S. helvola* populations had more consistently high within-population gene diversity (*H*_S_; expected heterozygosity; Nei, 1987), with the highest value in population IV-MO (0.125), and the lowest value again in population SNR-MO (0.086; Table 4). *S. leiosperma* populations IA and IV-MO had the lowest overall *H*_S_ at 0.078 and 0.076 respectively, while *S. leiosperma* population SNR-MO was higher at 0.101 (Table 4). Negative values for the inbreeding coefficient (*F*_IS_) were observed in *S. helvola* populations IA (−0.133) and SNR-MO (−0.019), and *S. leiosperma* populations IA (−0.387) and IV-MO (−0.310), indicating heterozygote excess (Table 4). *F*_IS_ was highest in *S. helvola* IV-MO, SP-MO, and AR (0.111-0.250), indicating heterozygote deficiency and possibly higher levels of inbreeding in these populations (Table 4). *S. leiosperma* population SNR-MO had an *F*_IS_ value of 0, indicating neither heterozygote deficiency nor excess, and thus approximate Hardy-Weinberg equilibrium (Table 4).

**Table 4.**
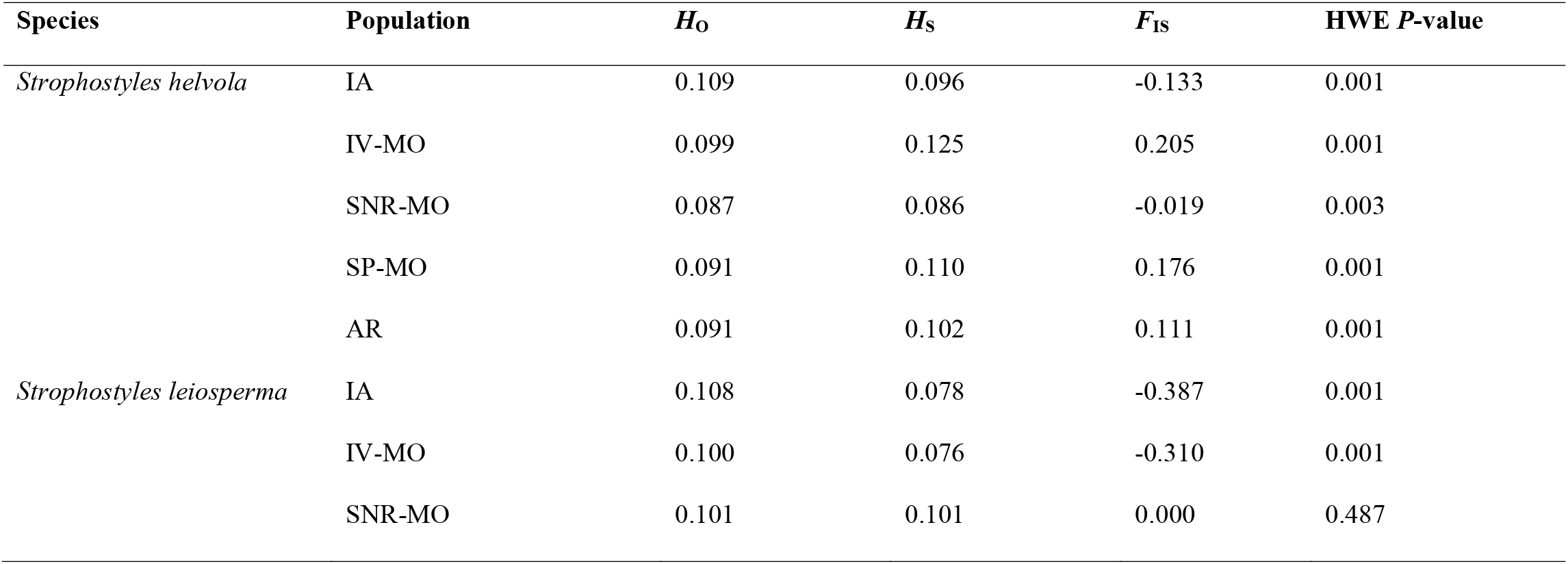
Diversity metrics for each population of *Strophostyles helvola* and *S. leiosperma. H*_O_ is the observed heterozygosity, *H*_S_ is within-population expected heterozygosity (Nei’s gene diversity), *F*_IS_ is the inbreeding coefficient (from an AMOVA), and the HWE *P*-value is for deviation of genotypes from Hardy-Weinberg equilibrium (random mating), where a significant value (*P* < 0.05) indicates significant deviation. *P*-values are derived from 1000 permutations and thus have a lower threshold of 0.001.

## DISCUSSION

We present the first population genetics study of *Strophostyles*, which supports the genetic separation of *S. helvola* and *S. leiosperma* and highlights unique patterns of genetic structuring and diversity among populations within each species.

### Separation of species

This study provides clear evidence for the genetic differentiation of *Strophostyles helvola* and *S. leiosperma*, at least in the range studied here. While these two species are sympatric across much of the central United States, there does not appear to be gene flow between them. This finding is consistent with the clear morphological distinction of *S. leiosperma* from the other two *Strophostyles* species, as well as genetic evidence from chloroplast and ITS markers (Riley-Hulting et al., 2004). This is in spite of similar pollen morphology across *Strophostyles* species, overlapping flowering times, generalist pollinators, and *S. helvola* and *S. leiosperma* being sister species (Krombein et al., 1979; Riley-Hulting et al., 2004; own observation). This clear genetic separation in spite of opportunity for hybridization indicates an as yet undetermined reproductive barrier between *S. helvola* and *S. leiosperma*. One possibility is interspecific pollen-rejection during pollen-pistil interaction (Bedinger et al., 2017); however, demonstrating this will require testing under controlled conditions.

### Population genetic structure within species

Population genetic structure was found to differ for *Strophostyles helvola* and *S. leiosperma. Strophostyles helvola* is a putatively outcrossing species with large, showy flowers and which is particularly common in mesic sites, while *S. leiosperma* is a putatively selfing species with small flowers and which tends to inhabit relatively more open, xeric sites; nevertheless, there is considerable overlap between their distributions (Fig. 1; Riley-Hulting et al., 2004). In *S. helvola*, we found four distinct genetic clusters among the five populations collected, with the two southernmost populations (SP-MO and AR) primarily assigning to the same cluster (Fig. 2). In *S. leiosperma*, we found only two genetic clusters, with the two northernmost populations assigning to the same cluster (Fig. 2).

Evidence of admixture was discovered primarily in three populations of *Strophostyles helvola*: IA, SP-MO, and AR, while the structure patterns of the other genetic clusters were largely unmixed. Given the great distances between collection sites (>200 km) and likely either selfing and/or insect-mediated pollination, it is unlikely that direct pollen flow is occurring across the populations studied (Hamrick, 1982; Riley-Hulting et al., 2004). Genetic admixture here could be from migration via seed dispersal and shared ancestry. Similar to the pattern in *S. helvola*, a study in *Capsella* found that in a selfing species, while there was genetic structure between populations, there were also several individuals in most populations with mixed genetic ancestry, which they attribute to the colonization of a few propagules from distant populations and subsequent genetic integration (St. Onge et al., 2011). However, there are also likely several populations of *S. helvola* and *S. leiosperma* interspersed between these localities, which may allow indirect gene flow between the populations studied here. Sampling from such interspersed populations at a finer geographic scale could reveal a genetic gradient of admixture produced by gene flow across these more proximate populations, which may thus function as metapopulations. Populations lacking admixture, such as *S. helvola* populations IV-MO and SNR-MO, may indicate that these populations are relatively isolated from such potential gene flow.

*Strophostyles helvola* may often be water-dispersed, as this species’ seeds are known to have a cellular coating which lends them buoyancy in water (Riley-Hulting et al., 2004). The seeds of *Strophostyles umbellata*, which have the same coating as *S. helvola*, have been observed to be buoyant in water for three weeks (Erickson and Young, 1995). The most proximate potential route of dispersal in our study region may therefore be the Mississippi River and its tributaries. However, while most of our sites are near tributaries of the Mississippi River, none of them lie directly downstream of sites at higher latitudes. Specifically, population IA is closest to the Cedar River (∼5 km); IV-MO is closest to the Des Moines River (∼3 km); SNR-MO is adjacent to the Meramec River (∼3 km); and AR is immediately adjacent to the Arkansas River. Population SP-MO is unique in that it is more than 16 km from the nearest river (Current River).

In contrast, *Strophostyles leiosperma* shows some population differentiation, but no substantive evidence of admixture (Fig. 2). The *S. leiosperma* population SNR-MO showed very high *F*_ST_ differentiation from *S. leiosperma* populations IA and IV-MO (Table 3). However, *S. leiosperma* samples from SNR-MO did not show qualitatively distinct floral or vegetative morphology compared to that of the other two populations. One explanation is that IA and IV-MO originated from the same ancestral source population or were connected via some route of migration. While *F*_IS_ in *S. leiosperma* population SNR-MO is 0, the reduced heterozygosity compared to the other two *S. leiosperma* populations may be due to a genetic bottleneck if the population was colonized by only a few founder individuals (Table 4). Unlike *S. helvola, S. leiosperma* seeds are generally smooth and not thought to be commonly water dispersed (Riley-Hulting et al., 2004). There are no currently recognized infraspecific taxa for *S. leiosperma*, although Riley-Hulting et al. (2004) did find a slight genetic difference (ITS region) between *S. leiosperma* samples from the northernmost / southwestern-most range and samples from Missouri and Arkansas.

Overall, the greatest amount of genetic variation for both species was found among rather than within populations (Table 2). However, this pattern could be primarily due to the large geographic distances between populations more so than a biological cause, as increasing distance between populations is known to decrease genetic diversity detected within populations and inflate diversity detected among populations (Reisch and Bernhardt-Römermann, 2014). Similarly, the great distances between populations may explain the lack of clear isolation by distance; at the scale of this study, local dispersal and gene flow may not be the predominant drivers of genetic differentiation (Twyford et al., 2020).

Human-mediated dispersal may also influence *Strophostyles* population dynamics, as evidenced by many herbarium records collected from anthropogenically disturbed sites, such as near railroads and ditches (Riley-Hulting et al., 2004). Riley-Hulting et al. (2004) specifically suggest *Strophostyles* seed dispersal via movement of ballast (gravel/sand track base) during the construction of railways (Riley-Hulting et al., 2004). *Strophostyles helvola* and *S. leiosperma* have repeatedly been observed in railroad ballast in the Midwest (Pepoon, 1927; Gilly and McDonald, 1936; Deam, 1940), likely due to their proclivity for sandy soil. Indeed, the municipalities of the sites SP-MO (Neelyville, Missouri) and AR (Little Rock, Arkansas) were directly connected by the St. Louis, Iron Mountain and Southern Railway as early as the 1870s (St. Louis, 1878; Stepenoff, 1993). In the absence of clear water routes, this may have been a conduit of shared ancestry in *S. helvola* between sites SP-MO and AR. Colonization of railway habitats has been observed in numerous plant species, particularly annual, ruderal species (Mühlenbach, 1979; Arnold, 1981; Austin, 2003; Hill and Blaney, 2009). The extent to which water and human dispersal has influenced population structure in *Strophostyles* could be more directly tested by sampling sites along major rivers and historic railroads. Rapid human dispersal could also contribute to nonlinear relationship between geographic and genetic distance.

### Population genetic diversity within species

Varied patterns of genetic diversity were also found within each species, which warrants further investigation. Heterozygote excess (negative *F*_IS_) was found in both *Strophostyles helvola* and *S. leiosperma*. High heterozygosity in *S. leiosperma* (IA and IV-MO; Table 4) is in contrast to the lack of heterozygosity found by Riley-Hulting et al. (2004) and the hypothesis that *S. leiosperma* is a primarily selfing species. However, Riley-Hulting et al. (2004) only sequenced two markers (trnK and ITS) and not from multiple individuals within populations, so they may have missed this signature of heterozygosity. Two populations of *Strophostyles helvola* also showed heterozygote excess (IA and SNR-MO; Table 4). Heterozygote excess can occur due to self-incompatibility, asexual reproduction, or selection for heterozygous individuals due to inbreeding depression (Stoeckel et al., 2006). This often occurs in longer-lived woody and clonal species (Stilwell et al., 2003; Ge et al., 2005; Stoeckel et al., 2006), although clonality and woodiness is not known to occur in *Strophostyles* (Riley-Hulting et al., 2004). On the other hand, there was also significant heterozygote deficiency among the other *S. helvola* populations IV-MO, SP-MO, and AR, which suggests some degree of inbreeding in these populations (Table 4). Mating system is known to be quite variable within plant species in general and can even vary among conspecific individuals within the same population (Hamrick, 1982; Ma et al., 2020). Such mating system variability may help explain the mixed patterns in genetic diversity in *Strophostyles*; testing outcrossing rate and self-compatibility across multiple populations will be needed to confirm this.

### Unique genetic patterns in population SNR-MO

At the Shaw Nature Reserve (SNR-MO), both *Strophostyles helvola* and *S. leiosperma* displayed unique patterns of genetic differentiation and diversity. Both species had distinctly differentiated genetic clusters for SNR-MO, with a lack of admixture with other populations (Fig. 2, 3), and SNR-MO consistently displayed the highest pairwise *F*_ST_ values with other populations (Table 3). Lastly, *F*_IS_ was close to 0 (Hardy-Weinberg equilibrium) in SNR-MO for both species (Table 4). All of this points to unique population dynamics occurring at SNR-MO, which could be related to the species’ unique history of establishment at this site. While no intentional introduction of *Strophostyles* is known at SNR-MO (James Trager, pers. comm. 2021), it is nevertheless possible that an undocumented or unintentional introduction occurred. Such an introduction may have involved only a small number of seeds, which could have induced a genetic bottleneck for both species. The lack of admixture with other populations also suggests that few if any further dispersal events occurred to introduce new genetic variation into this population, leaving it isolated. This possible founder effect is further suggested by the relatively lower level of heterozygosity for *S. leiosperma* in SNR-MO (*F*_IS_ = 0) compared to the other two populations (*F*_IS_ = −0.387 to −0.310; Table 4). The geographic origin of the propagules which colonized SNR-MO remains unknown, although the slight representation of the *S. helvola* SNR-MO genetic cluster in the southernmost populations SP-MO and AR suggest a potential origin from the southern range of the species (Fig. 2). Taken together, population SNR-MO serves as an interesting case study of the potential for both species to become isolated and genetically differentiate in certain disconnected populations. This warrants further investigation into the mechanisms responsible for genetic differentiation in this and other isolated populations, as well as whether differentiation was primarily produced by genetic drift or if there is an adaptive component.

### Future directions

Our study is limited in that a large portion of the range of both *S. helvola* and *S. leiosperma* could not be sampled. *Strophostyles leiosperma* extends across much of the central Great Plains region, and *S. helvola* also has a broad distribution across the coastal eastern U.S. (Riley-Hulting et al., 2004); novel genetic diversity likely remains to be discovered in those areas. Thus, it remains to be seen if our patterns of genetic differentiation and diversity apply to a broader sampling of both species, or if they are unique to the region studied here. Significantly high population differentiation found even at this scale suggests considerable genetic variation across the range of *Strophostyles* species. On the other hand, sampling from sites which are more geographically proximate will also be beneficial, in order to reveal further evidence of admixture if short-distance pollen transfer and seed dispersal are in fact important mechanisms for gene flow in these species. Ideally, future studies will examine *Strophostyles* genetic diversity across multiple geographic and biological scales. The alignment of both species to the *Phaseolus vulgaris* reference genome will also inherently constrain the amount of genetic variation that can be detected, which can be improved by constructing a de novo genome within the *Strophostyles* genus. Future work should also include the third species in the genus, *Strophostyles umbellata*, which has a separate distribution from the sampling in this study and is known to be strongly perennial (Riley-Hulting et al., 2004). There is also some morphological evidence for hybridization between *S. helvola* and *S. umbellata* (Riley-Hulting et al., 2004).

Overall, this study reinforces the need to further investigate multiple aspects of *Strophostyles* life history and population biology, particularly life span, mating system, and modes of dispersal, particularly whether these traits differ among populations studied and the ecological correlates of these differences. A precise study on the floral biology of *Strophostyles* species is also warranted, particularly on self-compatibility, interspecific pollen compatibility, anatomical differences in flower morphology, and pollinator interactions, which may all contribute to the observed patterns of genetic diversity and admixture.

## CONCLUSIONS

These data show for the first time the genomic diversity of *Strophostyles* species at the population level. From this, we were able to confirm the genetic separation of *S. helvola* and *S. leiosperma* within the range of this study, and we found complex patterns of genetic structure and diversity within both species. Mixed patterns of admixture and heterozygosity call for a detailed assessment of the life history and reproductive biology of both species, particularly mating system variation, as well as more fine-scale sampling allowing for clarification of local population dynamics. Information gained here demonstrates that there is an abundance of genetic diversity across the range of both *S. helvola* and *S. leiosperma*, and much of the distribution of these species remains to be explored. Acquiring genetic information from non-model taxa such as *Strophostyles* species will be critical in discovering novel genetic diversity related to environmental tolerances and other adaptive features that can inform plant evolutionary ecology and crop breeding.

## ACKNOWLEDGMENTS

This research was funded by the Perennial Agriculture Project (Malone Family Land Preservation Foundation and The Land Institute). S.A.H. was supported by a graduate research assistantship from the Saint Louis University Department of Biology. M.J.R. was supported by the Donald Danforth Plant Science Center and the Perennial Agriculture Project. We are grateful to the Department of Energy Joint Genome Institute and collaborators for pre-publication access to the *Phaseolus vulgaris* v.2.1 genome. We thank the following for their assistance in planning population sampling: Claudia Ciotir, Laura Klein, and Joel Swift. We are grateful to the following who assisted in locating collection sites: Steven Cannon, Laura Jackson, Justin Meissen, and Robert S. Wallace. For assistance with permitting and site information, we are indebted to Malissa Briggler, Christopher Crabtree, Ian Hope, Steve Paes, Daryl Smith, and James Trager. For assistance preparing samples and extracting DNA, we thank Niyati Bhakta, Claudia Ciotir, Emma Frawley, Nathan Held, Aidan Leckie-Harre, and Lisa Millar. We thank Donna Herrera at the Missouri Botanical Garden for assistance in depositing voucher specimens. We are grateful to the Carrington, Eveland, Nusinow, Slotkin, and Topp labs (Donald Danforth Plant Science Center) for providing necessary equipment and expertise in processing samples. We also wish to acknowledge the work of REU student Marissa C. Sandoval, for her additional studies on *Strophostyles* germination and herbarium morphological studies. Lastly, we appreciated the time and thoughtful feedback on previous versions of this manuscript from the Miller Lab and from researchers Wendy Applequist, Peter Bernhardt, Gerardo Camilo, Elizabeth Kellogg, Jason Knouft, and Chris Topp.

## AUTHOR CONTRIBUTIONS

S.A.H. and A.J.M. designed the study, and S.A.H. implemented the research and wrote the manuscript. Z.N.H. and M.J.R. assisted in the computational analysis of genomic data and contributed to the writing of the manuscript.

## DATA AVAILABILITY

All genomic data are deposited in the Sequence Read Archive, and R code is available at the following GitHub page. [Data is in preparation for submission to these permanent archives].

## SUPPORTING INFORMATION

Additional supporting information may be found online in the Supporting Information section at the end of the article.

**APPENDIX S1**. Excel spreadsheet containing metadata for all samples collected, including location coordinates and details, and genetic quality metrics.

**APPENDIX S2**. Genetic structure plots with K=2 to K=7 clusters for *Strophostyles helvola* and *S. leiosperma*.

## Notes

### Competing Interest Statement

The authors have declared no competing interest.

