## Appendix S2 for "Comparative population genomics in two sympatric species of *Strophostyles* (Fabaceae) with contrasting life histories"

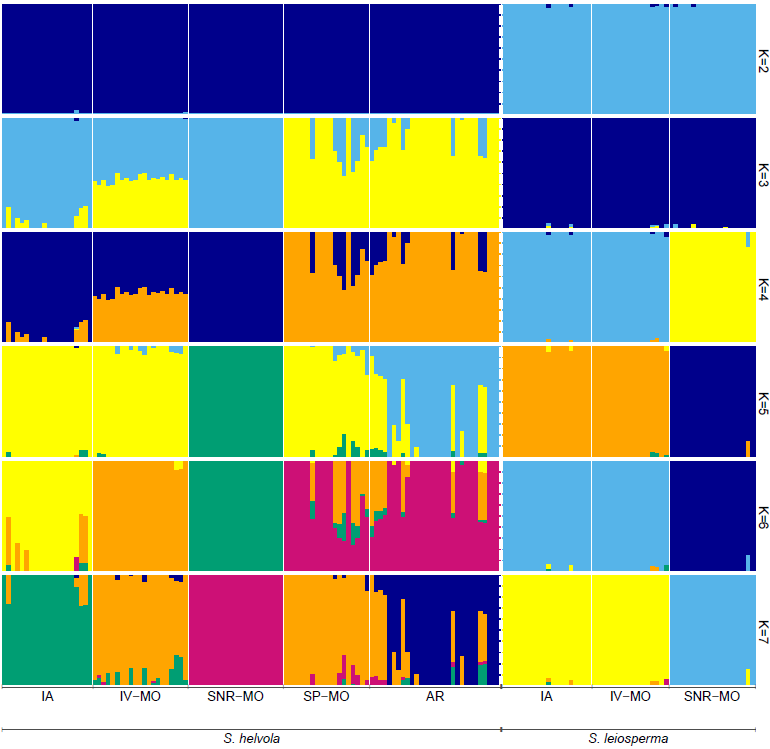

**Appendix S2.** Genetic structure plots from K=2 to K=7 clusters for *Strophostyles helvola* and *S. leiosperma*; K=7 clusters produced model complexity that maximized marginal likelihood, although the seventh cluster has an extremely low representation, and thus effectively only six clusters are visible. Populations, labeled below the bars, are ordered north to south within species (left to right) and are separated by thin white solid lines; species are separated by a thick white dashed line. Each vertical bar represents a single individual from each population. Each color signifies a unique genetic cluster assignment, with multiple colors within an individual signifying mixed ancestry.
